# A measure of concurrent neural firing activity based on mutual information

**DOI:** 10.1101/2020.12.16.423134

**Authors:** Gorana Mijatovic, Tatjana Loncar-Turukalo, Nebojsa Bozanic, Luca Faes

## Abstract

Multiple methods have been developed in an attempt to quantify stimulus-induced neural coordination and to understand internal coordination of neuronal responses by examining the synchronization phenomena in neural discharge patterns. In this work we propose a novel approach to estimate the degree of concomitant firing between two neural units, based on a modified form of mutual information (MI) applied to a two-state representation of the firing activity. The binary profile of each single unit unfolds its discharge activity in time by decomposition into the state of neural quiescence/low activity and state of moderate firing/bursting. Then, the MI computed between the two binary streams is normalized by their minimum entropy and is taken as positive or negative depending on the prevalence of identical or opposite concomitant states. The resulting measure, denoted as Concurrent Firing Index based on MI (CFI_MI_), relies on a single input parameter and is otherwise assumption-free and symmetric. Exhaustive validation was carried out through controlled experiments in three simulation scenarios, showing that CFI_MI_ is independent on firing rate and recording duration, and is sensitive to correlated and anti-correlated firing patterns. Its ability to detect non-correlated activity was assessed using ad-hoc surrogate data. Moreover, the evaluation of CFI_MI_ on experimental recordings of spiking activity in retinal ganglion cells brought insights into the changes of neural synchrony over time. The proposed measure offers a novel perspective on the estimation of neural synchrony, providing information on the co-occurrence of firing states in the two analyzed trains over longer temporal scales compared to existing measures.

## 1 Introduction

The role of synchrony in nervous system signaling has been analyzed in many different ways, mostly as the key indicator of a transmission of temporally precise action potentials - spikes (Cutts and Eglen, 2014). Quantifying the level of synchronized firing activity in inherently noisy environments is indeed of utmost importance for the general understanding of both stimulus-induced and internal coordination of distributed neuronal responses (Singer, 1999; Ermentrout et al., 2008; Grewe et al., 2017).

Numerous studies have investigated correlated neural activity in early sensory pathways, contributing to the question of processing and decoding of relevant sensory information in the brain. It has been shown that firing patterns in the optic nerve are strongly shaped by synchronized activities in the inner retina (Brivanlou et al., 1998). Concerted firings from many retinal ganglion cells (RGCs) could provide subtle visual information about the presence of fine spatial details (Meister et al., 1995; Schnitzer and Meister, 2003), or of information redundancy related to visual features that mainly originate from retinal neighbors (Puchalla et al., 2005; Schneidman et al., 2003). Synchronized relative spike timings convey information about visual stimuli (Gollisch and Meister, 2008), and a strong impact of correlated spike trains on the formation of retinotopic maps has been found (Xu et al., 2011). In addition, synchronous firing of ensembles of neurons has been found in the visual pathway, analyzing more aspects such as the perceptual consequences of synchronous activity, the ability to carry information, and the transmission of synchronous neural activities to subsequent stages of processing (Usrey and Reid, 1999). It has been revealed that visual synchrony activities affect the ability of grouping and segmentation in perception (Usher and Donnelly, 1998). Recently there has been also evidence on the relevance of synchronous activity for obtaining a synchrony code, although other features operating at the single unit level are important for a complete encoding of time-dependant stimuli (Grewe et al., 2017). All these works provide evidence that understanding the link between correlated activity and the representation of sensory information yields insight into the characterization of neural interactions in general (Brown et al., 2004), and also brings into question how these interactions influence the processing of information in higher cortical areas (Buzsaki, 2006; Kandel et al., 2000).

A wide variety of measures have been developed for estimating the degree of synchronous activity between pairs of neurons. The terminology convention has not been clearly established and measures are usually named after the underlying methodology, defining different indices of neural synchrony. Thirty-five pairwise measures are listed in a review paper (Cutts and Eglen, 2014), which includes various approaches to detect pairwise correlation in spike trains and provides a systematical comparison among them. Some categories comprise: measures quantifying the distance between spikes trains relying on instantaneous (dis)similarities, as in the case of time-resolved ISI-distance and SPIKE-distance indices (Kreuz et al., 2015); measures counting pairs of spikes which are neighbors of each other (Wong et al., 1993; Pasquale et al., 2008); information-theory based measures (Gray, 2011; Li, 1990); measures modeling spike trains as a marked point processes (Schlather et al., 2004). The authors of (Cutts and Eglen, 2014) identified the necessary and desirable properties needed for a correct quantification of correlated spiking activities. The necessary properties include: symmetry, robustness to variations in firing rate (FR) and recordings’ duration, a bounded range in [−1, 1] for clear distinction of identical, anti-correlated and uncorrelated firing activity, and robustness to small variations in a commonly used parameter - the window of synchrony. The desirable properties involve avoiding counting quiescence (inactive) periods as correlated, a minimal number of input parameters, and minimal or no assumptions on the underlying ISI distribution (Cutts and Eglen, 2014). Out of the measures reviewed, three of them are reported to satisfy all the necessary properties: the Spike Count Correlation Coefficient (SCCC) (Eggermont, 2010), its improved version denoted as Kerschensteiner and Wong correlation (KWC) (Kerschensteiner and Wong, 2008), and the Spike Time Tiling Coefficient (STTC) (Cutts and Eglen, 2014).

The large majority of the proposed synchrony measures quantify the degree of association between simultaneously recorded streams through a direct comparison of time-aligned consecutive spike timings or inter-spike intervals in the two streams. This approach allows to develop measures that are well-resolved in time and able to capture synchrony over short time scales usually defined by a window of synchrony, *Δt*. However, it is also exposed to issues such as the dependence of the synchrony measures on the pre-defined temporal scale (related to *Δt*), and the sensitivity to both measurement errors (e.g., errors in spike sorting) and physiological errors (e.g., random jitters in the spiking times of synchronized neurons). All these issues are exacerbated by the need of capturing the fine temporal structure of neural spike trains and of incorporating it into the synchrony measure.

In this work we undertake a different approach, inspired by the field of joint symbolic dynamics (Porta et al., 2015) and the characterization of firing patterns introduced in (Mijatović et al., 2018): we deliberately discard the fine structure of spiking times in the two analyzed neuronal units, and rather perform a coarse characterization of their firing dynamics tailored to identify states of activity and inactivity for each neural unit. The coarse state representation further enables pairwise estimation of the co-occurrence of firing states in the two trains, so as to reflect neural synchrony over longer temporal scales. Such estimation relies on a modified measure of mutual information (MI) applied to the binary representation of the spiking activity unfolded in time. The resulting Concurrent Firing Index based on MI, CFI_MI_, is assumption-free, requires only one input parameter, is symmetric and bounded between −1 and 1, and can distinguish uncorrelated, correlated and anti-correlated concomitant firing patterns. In a preliminary research published in a conference paper (Mijatovic et al., 2020, accepted paper), the CFI_MI_ index was compared with SCCC, KWC and STTC indices in controlled experiments of synthetic Poisson spike trains. In this work, we further exploit Poisson spike trains to validate thoroughly the robustness of the index to variations in firing rate and recording duration, also assessing its dependence on its free input parameter. Moreover, we explore the informative value of CFI_MI_ in the assessment of pairwise correlations in a more complex simulated network scenarios, and more importantly on experimental recordings of retinal ganglion cells.

## 2 Materials and Methods

### 2.1 Discrete state representation of firing activity

The activity over time of a single neural unit is commonly represented as a spike train, i.e. a series of action potentials (AP) precise timing. As a time series of discrete events, a spike train can be considered as a temporal point process with periods of intensive and moderate firing, intertwined with quiescent intervals (Brown et al., 2004). Consequentially, intervals between consecutive spikes, inter-spike intervals (ISIs), can be coarsely grouped into three classes: very short ISIs, ISIs of moderate duration and long ISIs. This concept of firing representation exploiting the information on ISI duration and classifying ISIs into three states unfolded in time, represents the basic probabilistic approach for a description of the firing patterns originally proposed in (Mijatović et al., 2018). According to this approach, three functional states are defined, as determined by two pre-specified thresholds: 1) bursting state (**B**) − including all ISIs shorther than the bursting threshold, *thr*_**B**_ (very short ISIs), 2) idle state (**I**) − including all ISIs longer than the idle threshold, *thr*_**I**_ (very long ISIs) and 3) firing state (**F**) − including all ISIs between these two thresholds (i.e. ISIs of moderate duration) (Mijatović et al., 2018).

The decision boundary between the states **B** and **F** is determined by the bursting threshold *thr*_**B**_, specified as the minimum ISI below which the dense firing pattern can be characterized as *bursting* activity, i.e. activity exhibiting very short ISIs between consecutive spikes (Izhikevich, 2000). Thus, the threshold *thr*_**B**_ represents a physiological constant which characterizes the cell’s bursting ability and depends on the brain region (Mizuseki et al., 2012). The quiescent ISI intervals reflect periods of very low or no activity and should be longer than an average inter-spike interval. In (Mijatović et al., 2018), ISI intervals are considered quiescent if equal to or longer than a threshold *thr*_**I**_, being thus classified in the idle state **I**. The estimation of the idle threshold is directly related to the intrinsic firing properties of each neuron individually, i.e. to the statistical properties of its entire ISI stream, denoted as **ISI**. This threshold is estimated as: *thr*_**I**_ = *b* · *mean*(**ISI**); where *mean*(**ISI**) is the average ISI duration of **ISI**, while *b* is an adaptive parameter for the detection of quiescence periods. The idle threshold estimation and the selection of *b* have been extensively investigated in (Mijatović et al., 2018). Between the bursting and idle state, ISIs longer than *thr*_**B**_ and shorter than *thr*_**I**_ are considered as moderate firing and classified into the **F** functional state.

The three-mode decomposition of neural firing patterns served in this study as a starting point to formulate a measure of concurrent firing activity between pairs of neurons. The first step provides a binary representation of the spiking activity of each neural unit, distinguishing firing and non-firing periods throughout the recording time. To do this, the **F** and **B** functional states are integrated into one single working state (**W**, binary encoded as **1**), and the remaining state **I** is labeled as non-working state (**nW**, binary encoded as **0**). The resulting binary flow contains information on firing (**W**) and non-firing (**nW**) periods distinguished by the idle threshold *thr*_**I**_. The spiking activity decomposition into **B, I** and **F** states for two exemplary spike trains is illustrated in Fig. 1 together with a state-diagram of the **W** and **nW** states and the corresponding binary profiles.

**Fig. 1.**
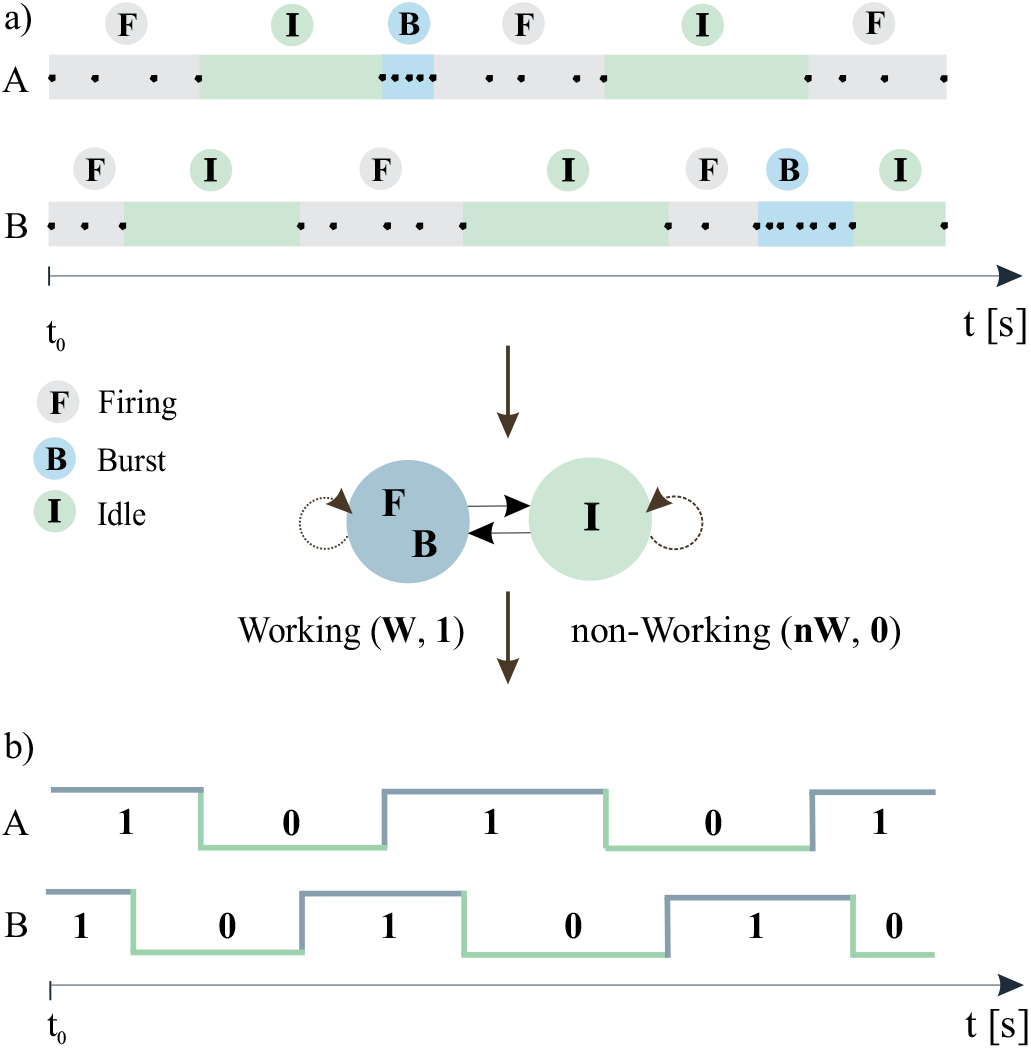
Construction of binary streams from two spike trains. a) Two exemplary trains A and B with firing patterns decomposed into three functional states (**F, B, I**), and binary state-diagram obtained merging the states **F** and **B** into a single working state (**W, 1**), while **I** represents the non-working state (**nW, 0**); b) the corresponding binary profiles of trains A and B as a function of time.

### 2.2 Concurrent firing index based on mutual information

Mutual information (MI) is a symmetric, nonlinear measure of dependence between two random variables, well-known in information theory (Shannon, 1948). MI can be interpreted as the amount of information shared by the two variables, or as the degree of predictability provided by one variable about the other one. In this paper, we use MI to quantify the dependence between the binary spiking profiles of the two considered trains, obtained as explained in Sect. 2.1. Moreover, we modify it to obtain a measure which is normalized and can reflect both correlated and anti-correlated spiking activity.

In order to compute MI between the profiles of two spike trains, encoded in the two random variables A and B, first the joint and marginal probabilities of A and B are quantified as follows:

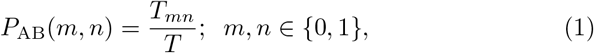

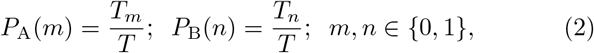

where *T*_*m*_ is the time during which the train A is in the state *m, T*_*n*_ is the time during which the train B is in the state *n, T*_*mn*_ is the time during which the train A is in the state *m* while the train B is in the state *n*, and *T* is the total time of recording. Then, the MI between A and B is defined as (Shannon, 1948):

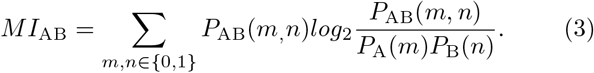

*MI*_AB_ is bounded between zero and the minimum entropy of the two variables A and B, *H*_*min*_ = *min*{*H*_A_, *H*_B_} (Gray and Shields, 1977), where the entropy of A is computed as:

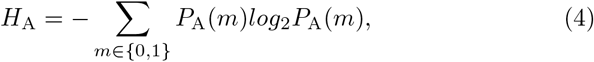

and the entropy of B is computed analogously. To get a normalized measure, *MI*_AB_ is adjusted dividing it by *H*_*min*_, so that to reach a maximum absolute value of 1, corresponding to fully dependent variables A and B. Moreover, an additional modification is performed to make the index able to reflect correlations and anti-correlations. This is achieved by assigning to the normalized MI measure a positive or negative sign on the basis of the tendency of the two variables to be in the same state (indicative of positive correlation) or in opposite states (indicative of negative correlation). Specifically, we compute the probability *p*_*c*_ as the average conditional probability that train A is in a certain state given that train B is in the same state, and the probability *p*_*ac*_ as the average conditional probability that train A is in a certain state given that train B is in the opposite state:

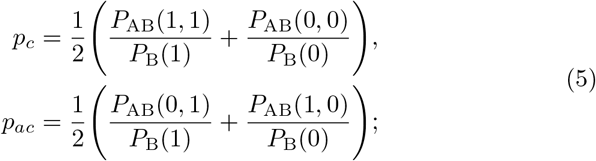

then, we assign positive synchrony values to the case in which the co-occurrence of the same state for A and B is more frequent (*p*_*c*_ *> p*_*ac*_) and negative values to the case in which the occurrence of different states for A and B is more frequent (*p*_*ac*_ *> p*_*c*_). Accordingly, the proposed concurrent firing index based on mutual information, denoted as CFI_MI_ index, is defined as:

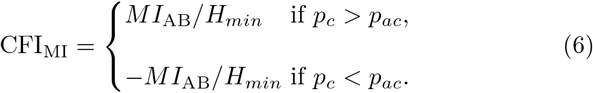

Note that when the two variables A and B are statistically independent the two probabilities in eq. (5) are the same (*p*_*c*_ = *p*_*ac*_ = 0.5) and the concurrent firing index vanishes (CFI_MI_ = 0). We cover also the cases in which one or both the trains stay constantly in the same functional state (**W**/**nW**) over time; in such cases, for which eq. (6) is undefined as *H*_*min*_ = 0, we assign CFI_MI_ = 0 if only one train is steady over time, CFI_MI_ = 1 if both trains are steadily in the same state, and CFI_MI_ = −1 if both trains are steadily in opposite states.

The CFI_MI_ measure inherits the properties of MI, being symmetric and sensitive to both linear and nonlinear correlations between the discretized spiking activity of two trains. Moreover, it spans the bounded range [−1, 1] commonly used to reflect correlations and anti-correlations, being equal to 1 in case of fully correlated trains, equal to zero in case of uncorrelated trains, and equal to −1 in case of fully anti-correlated trains. These properties are illustrated in Fig. 2 in a theoretical example that reproduces varying conditions of coupling between two binary random variables representing the discretized activity of two spike trains. In such example, we set the marginal probabilities of the trains A and B to be in working mode at the fixed values *P*_A_(1) = *P*_B_(1) = 0.5, and vary the joint probability of A and B to be simultaneously in working mode, *p* = *P*_AB_(1, 1), in the range *p* ∈ [0, 0.5]. From these values, all joint and marginal probabilities can be computed analytically for each value of *p* exploiting the marginalization property (Shannon, 1948), finding *P*_AB_(1, 1) = *P*_AB_(0, 0) = *p* and *P*_AB_(0, 1) = *P*_AB_(1, 0) = 0.5 − *p*; all quantities in eqs. (3 – 6) can be derived accordingly. The condition *p* = 0 corresponds to fully anti-correlated variables (Fig. 2b, case (1)), for which *MI*_AB_ = 1 with *p*_*c*_ = 0, *p*_*ac*_ = 1, so that the proposed index takes the value CFI_MI_ = −1. Increasing the probability *p* decorrelates the two variables because their joint entropy grows with constant marginal entropies, up to the value *p* = 0.25 for which A and B are independent (*P*_AB_(*m, n*) = *P*_A_(*m*)*P*_B_(*n*), Fig. 2b, case (2)) and thus *MI*_AB_ = CFI_MI_ = 0 (in this case, *p*_*c*_ = *p*_*ac*_ = 0.5). Increasing the parameter *p* above 0.25 makes the variables positively correlated leading to positive values of CFI_MI_ (*p*_*c*_ *> p*_*ac*_), up to the condition of maximum positive correlation attained for *p* = 0.5 (Fig. 2b, case (3)) and documented by the maximum information shared between A and B (*p*_*c*_ = 1, *p*_*ac*_ = 0, *MI*_AB_ = CFI_MI_ = 1).

**Fig. 2.**
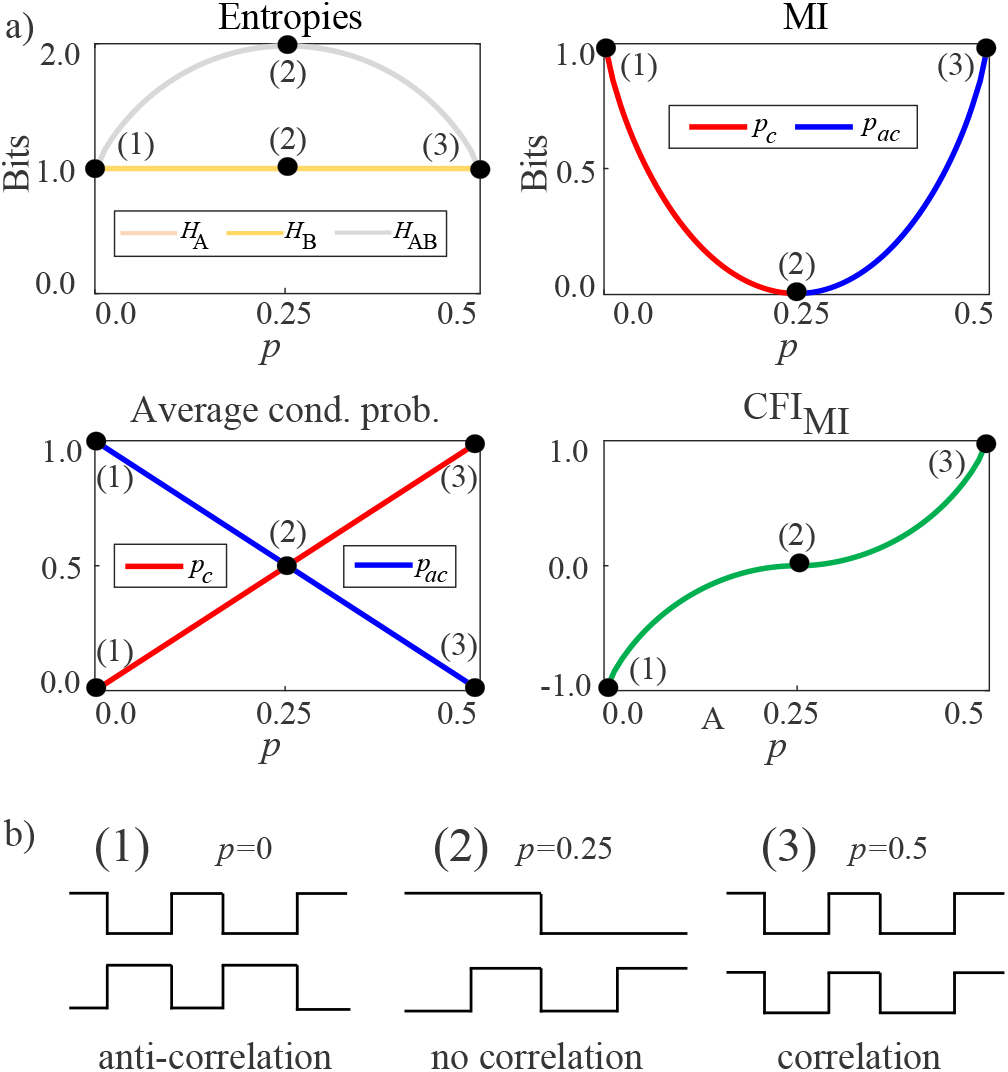
Computation of the CFI_MI_ index in a theoretical example involving two binary random variables A and B. Information-theoretic functionals leading to CFI_MI_, computed as a function of the probability *p* that the variables A and B are both equal to 1; b) exemplary profiles of A and B for three representative values of *p*.

### 2.3 Simulations

In this section, we present the three simulation scenarios used to test and evaluate the behavior of the proposed index with respect to the necessary properties of neural correlation measures. The first and second scenarios include respectively independent Poisson spike trains and mutually dependent spike trains coupled to reproduce both correlated and anti-correlated dynamics. The third simulation assumes a realistic networked scenario with coupled cortical dynamics (Izhikevich, 2003). In the simulations we test also the dependence of the proposed index on its only input parameter, i.e. the parameter *b* used for determination of the idle threshold (i.e. non-working mode); in all experiments, *b* is changed in the range {1, 2, 3, 4, 5}.

#### 2.3.1 Independent Poisson spike trains

The robustness of the CFI_MI_ index on firing rate (FR) and recording time duration was tested generating independent spike trains with Poisson distribution. For both experiments, 100 pairs of independent Poisson spike trains were simulated. When robustness to the changes in FR was tested, one train was generated with fixed FR = 1 spike/s, while the FR of the other train was varied in the range [1, 10] with a step of 1 spike/s, keeping the recording time *T* fixed to 300 s. To examine the dependence on the recording time, both trains were simulated with fixed FR = 3 spikes/s and *T* was varied in the range [30, 50, 100, 200, 300, 500, 1000] seconds.

#### 2.3.2 Dependent Poisson spike trains

The performance of the CFI_MI_ index in quantifying the degree of synchrony of coupled spike trains was evaluated in a second simulation in which both correlated and anti-correlated dynamics can be obtained. The designed spike train generator produces two trains such that the firing rate of one spike train is modulated locally in time depending on the firing of a master spike train. The master spike train A was generated as Poisson train with fixed FR = 3 spikes/s for a total duration of *T* = 300 s. After counting the total number of spikes in the master train (*N*_S_) and computing the average ISI duration 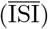, periods of ‘fast’ and ‘slow’ spiking activity, including ISIs shorter or longer than a limit threshold 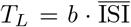, were identified in train A, and the total duration of these periods, *T*_*f*_ and *T*_*s*_ was determined (*T* = *T*_*f*_ + *T*_*s*_). Fig. 3a) illustrates an exemplary master train with ISIs belonging to *T*_*f*_ and *T*_*s*_ overlined with red and blue, respectively. Then, the spike train B was generated based on the master spike train A using a random spike insertion modulated by the parameter *γ*. Specifically, for each ISI in train A, a certain number of spikes was inserted randomly in the same time interval in train B using the following rule: if *ISI < T*_L_, a number of spikes equal to *γ* · *N*_S_ · *ISI/T*_*f*_ was inserted; if *ISI >*= *T*_L_, (1 − *γ*) · *N*_S_ · *ISI/T*_*s*_ spikes were inserted. The train B generated in this way contains the same number of spikes as the train A, with allocation depending on parameter *γ*. The parameter *γ* was varied between the extremes *γ* = 0, yielding a train B whose working periods coincide with the non-working periods of the master train A, and *γ* = 1, yielding a train B whose working periods coincide with the working periods in A. Fig. 3b) reports exemplary simulations of anti-correlated (*γ* = 0) and correlated (*γ* = 1) spike train activity.

**Fig. 3.**
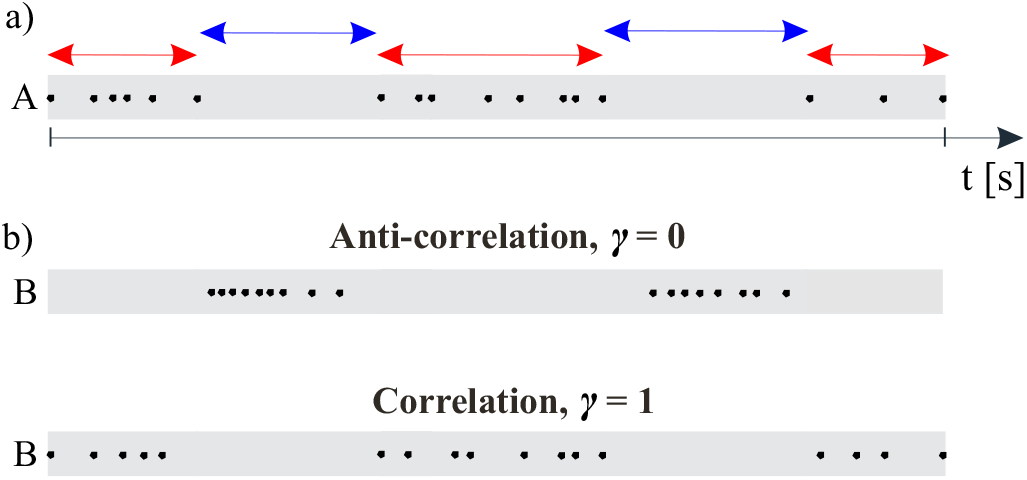
Exemplary spike trains generated in the second simulation. a) Master spike train A; b) spike train B generated to be anti-correlated with A, *γ* = 0; c) spike train B generated to be correlated with A, *γ* = 1. Red and blue arrows overline the duration of ISIs shorter and longer than the limit threshold (*T*_*L*_), respectively.

#### 2.3.3 Realistic spiking model for producing coupled cortical dynamics

To test the ability of the CFI_MI_ index to capture different levels of coupling in interconnected neurons similar to those of the mammalian cortex, we used the model proposed in (Izhikevich, 2003). This simple network model relies on well-established models for initiation and propagation of APs in neurons: the Hodgkin-Huxley model which ensures biologically plausible dynamics, and the quadratic integrate-and-fire model which guarantees computational efficiency. The overall model belongs to a class of pulse-coupled neural networks, where the neurons receive synaptic and noisy thalamic inputs. It is based on two differential equations and uses four parameters, whose variation allows to produce the dynamics of several fundamental cell types observed in neocortical neurons of mammals (Izhikevich, 2007).

Here, we simulate excitatory cells with regular spiking (RS) and inhibitory cells with low-threshold spiking (LTS), setting the ratio of excitatory to inhibitory neurons in the network to 4:1 for reproducing the mammal’s cortex (Izhikevich, 2003). We implemented a large-scale RS-LTS neural network with preserved neural heterogeneity, consisting of 1000 randomly connected neurons, with 800 RS excitatory cells and 200 LTS inhibitory cells. The network generation produces a random synaptic matrix *S*, whose elements *s*_*ij*_ have values in the range [−1, 1] reflecting the strengths of the synaptic connection between the output of the *j*^*th*^ neuron and input to the *i*^*th*^ neuron (*i, j* = 1, …, 1000). This initially generated synaptic matrix *S* was considered to reflect the nominal synaptic strengths, and was stored for future reuse to preserve the initial settings. Then, the RS-LTS networks are simulated several times more, each time using a scaled synaptic matrix, *S*_*α*_ = *α* · *S*, to preserve the original connections and uniformly modify their strengths. The scaling coefficient *α* was decreased from 1 to 0.2 in steps of 0.2. The total duration of each simulation was set to 30 s.

### 2.4 Experimental data

To identify concomitant firing patterns in a real set of neural responses, we analyzed experimental recordings acquired across the ganglion cell layer (GCL) in mouse retinal explants (Milosavljevic et al., 2018). In the following we briefly explain the motivation and design of experimental protocol in (Milosavljevic et al., 2018). The rate of information transfer through the optical nerve relies on the spiking activity from the inner retina which encodes the visual stimulus in an energetically optimized way (Spavieri et al., 2010). As the illumination of the ambient scene increases, the noise decreases and more information can be extracted from the visual scene (Milosavljevic et al., 2018). There is a specific class of retinal ganglion cells (RGCs) optimized for encoding the illumination of the visual scene, known as intrinsically photoreceptive retinal ganglion cells (ipRGCs) (Brown et al., 2010; Allen et al., 2017; Wong, 2012), which potentially serve as modulators of RGCs activity facilitating efficient visual information transfer at higher illumination levels (Milosavljevic et al., 2018). In order to test the hypothesis that firing rates for RGCs adjust with illumination level sensed by ipRGCs, in (Milosavljevic et al., 2018) a multielectrode array (MEA) extracellular recording was performed across the GCL of six mouse retinal explants exposed to a repeated sequence of temporal white noise (WN) added on a gradual illumination ramp in the absence of any other visual stimulus. It is worth noting that the WN amplitude was scaled with the illumination level. The gradual illumination ramp served to induce a response to slow changes in ambient light, whereas the superimposed WN simulated the high frequency visual stimulus. The total stimulus presentation in the dataset provided by the authors is 900 s, during which the illumination was gradually increased from 11.8 to 14.8 log10 photons · cm ^*−*2^ · s −1. Additionally, we have used the provided information on differentiation of RGC cells with respect to function into neurons with ON and OFF-type response to increments in radiance. To identify ON and OFF cells among 790 RGCs extracted from 6 retinas, the spike-triggered averages (STAs) were computed for WN stimulus (Milosavljevic et al., 2018) and the corresponding labels were provided with a dataset. From 790 cells available, 709 RGCs have produced statistically significant STA comprising: 406 ON-type response cells and OFF-type cells (303) (Milosavljevic et al., 2018).

All other details of the experimental protocol including ethical approval are provided in (Milosavljevic et al., 2018). The original study has included two more experimental scenarios to test the hypotheses regarding a mechanism of proactive control of information transfer capacity in retinal circuitry. However, this study uses only the described scenario, as our motivation was not the reproduction or validation of published results, but rather the evaluation of the proposed measure as regards its potential to uncover patterns of concomitant activation among RGCs in realistic settings, with inherent problems brought by measurement noise and pre-processing procedures (e.g., spike-sorting).

### 2.5 Assessment of statistical significance

The statistical significance of the synchrony values computed in simulated and experimental data was assessed, in the context of statistical hypothesis testing, computing the CFI_MI_ index for the analyzed pair of spike trains and over a set of surrogate pairs generated under the null hypothesis of uncorrelated trains. Surrogate spike trains were generated using a recently proposed method specifically designed for point processes, denoted as JOint DIstribution of successive inter-event intervals (JODI)(Ricci et al., 2019). The JODI algorithm generates, in a reliable and computationally efficient way, surrogate data retaining the same amplitude distribution and approximating the auto-correlation of the original inter-event intervals, while destroying any coupling. As in standard approach for statistical significance assessment, we compared the CFI_MI_ index computed for a given pair of spike trains with the distribution of CFI_MI_ assessed over 100 pairs of surrogate spike trains obtained through repeated application of the JODI algorithm. Then, according to a two-tailed hypothesis test with 5% significance, the CFI_MI_ index was regarded as indicative of: statistically significant anti-correlation between the two original spike trains if its value was lower than the 2.5^*th*^ percentile of its distribution on the surrogates; absence of correlation if its value was between the 2.5^*th*^ and 97.5^*th*^ percentiles of the surrogate distribution; or positive correlation if its value was higher than the 97.5^*th*^ percentile of the surrogate distribution.

## 3 Results

### 3.1 CFI_MI_ performance on simulated data

The proposed CFI_MI_ index was first evaluated on pairs of independent Poisson spike trains. The analysis was performed to assess the robustness of CFI_MI_ to variations in the input parameter, in the FR ratio between the two considered spike trains, and in the duration of the recordings, as specified in Sect. 2.3.1. For each analysis setting, 100 realizations of independent Poisson trains were generated and, for each realization, the statistical significance of CFI_MI_ was assessed using the JODI surrogates s reported in Sect. 2.5. The results in Fig. 4 document very stable estimates of the index in all settings, as the CFI_MI_ values were always distributed around zero while increasing the parameter *b* from 1 to 5 (Fig. 4a), while increasing the FR ratio from 1 to 10 (Fig. 4b), or while changing the recording time from 30 to 1000 sec (Fig. 4c). The variance of the estimates showed a tendency to increase for high values of *b*, while decreasing the FR ratio, and as expected decreasing the recording time. In all cases, the number of false positive couplings was around the nominal 5 % significance set for hypothesis testing, thus demonstrating the ability of the proposed measure to reflect the absence of correlated spike train activity with non significant values of CFI_MI_.

**Fig. 4.**
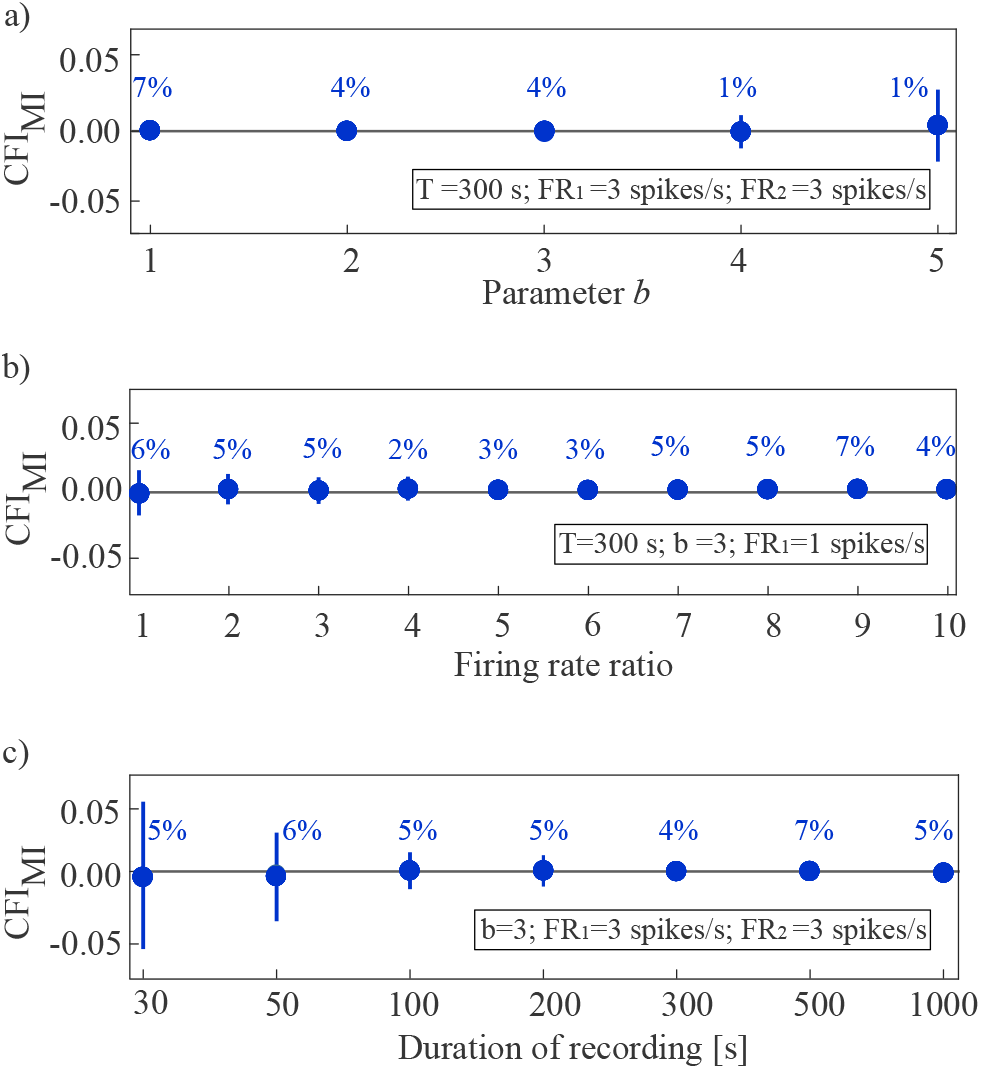
Simulation of independent Poisson spike trains. Distributions of the CFI_MI_ index (mean ± standard deviation) across 100 realizations of independent Poisson spike trains estimated under different conditions by varying only: a) the input parameter *b* used for detection of the idle threshold reflecting periods of neuron’s quiescence; b) the ratio of between the firing rates of the two spike trains; and c) the duration of the simulated recordings. The numbers above the index values, colored in blue, indicate the percentage of independent spike trains realizations for which the null hypothesis of independence was rejected.

Second, we studied the CFI_MI_ index for the simulation scenario of dependent Poisson spike trains introduced in Sect. 2.3.2. Simulations were performed varying the simulation parameter *γ* from 0 to 1 in steps of 0.05, in order to study how the index reflects coupling conditions ranging from anti-correlation to correlation. Moreover, the analysis was repeated for different values of the input parameter in the set *b* ∈ {1, 2, 3, 4, 5}, to investigate the dependence of the detected coupling profiles on the threshold that separates the working and non-working modes in the binary representation of each spike train. For each combination of *γ* and *b*, 100 realizations of the simulation were generated, and for each realization the statistical significance of the CFI_MI_ index was assessed through surrogate data analysis. The results reported in Fig. 5 depict the distribution of the index (a) and the percentages of statistically significant values (b). We found that, regardless of the choice of the input parameter, the proposed measure can distinguish correlated from anti-correlated cases. Indeed, for each value of *b*, the CFI_MI_ distribution moves from negative to positive values when *γ* increases from 0 to 1 (Fig. 5a) and the statistical significance analysis shift from full detection of negative correlations to full detection of positive correlations (Fig. 5b).

**Fig. 5.**
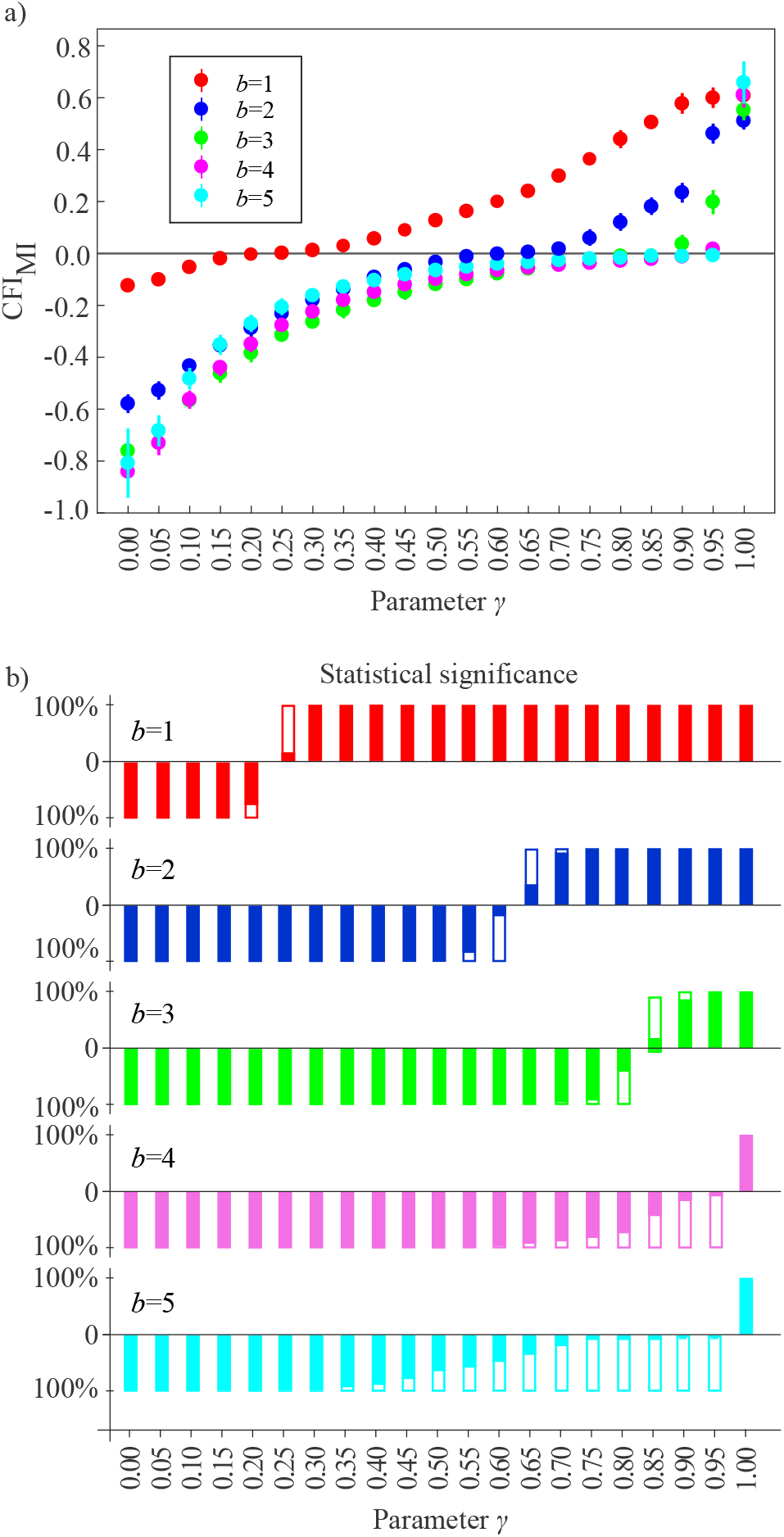
Simulation of dependent Poisson spike trains. a) Distribution of the CFI_MI_ index (mean ± standard deviation over 100 realizations of the simulation) computed as a function of the coupling parameter *γ* for different values of the input parameter *b*; b) percentage of realizations exhibiting significant positive correlation (filled bars above zero), absence of correlation (unfilled bars) or significant anti-correlation (filled bars below zero).

However, the change in the sign of the CFI_MI_ estimates and the switch from negative to positive correlations occur for higher values of *γ* when *b* increased from 1 to 5. This behavior is likely related to the fact that the parameter *b* sets the definition of the non-working mode: *b* = 1 is an extreme case where any ISI longer than the mean ISI is assigned to periods of inactivity, whereas case *b* = 5 assumes that only 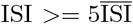 are classified as periods of quiescence. It is also noticeable that increasing *b* enlarges the range of *γ* values for which spike trains dominantly exhibit absence of correlation; for example, for *b* = 5 the range of values *γ* ∈ [0.55, 0.95] indicates absence of correlation. This result suggests that lower values of *b* are more appropriate to allow sharp transitions in the detection of anti-correlated vs. correlated spiking activities.

The third simulated scenario is relevant to the realistic network model with 1000 coupled neurons described in Sect. 2.3.3. For computational reasons, and without loss of generality because of the homogeneity of the network, we randomly selected RS (16) and LTS (4) units, and evaluated the changes in the CFI_MI_ index due to a gradual weakening of the synaptic strength (coefficient *α* of the synaptic matrix decreasing from 1 to 0). The randomly selected RS and LS units were split in two halves, maintaining the proportion between excitatory and inhibitory units, and the resulting 10 × 10 sub-network was considered for the computation of the CFI_MI_ index. The distribution of the CFI_MI_ estimates obtained for these 100 pairs of neural responses is presented in Fig. 6a), while Fig. 6b) presents the corresponding percentage of pairs detected as significantly correlated according to surrogate data analysis. The analysis is performed considering different values of the scaling coefficient *α* and computing the index with different values of the input parameter *b*. As expected, the estimated coupling was strong and statistically significant for the simulated spike trains obtained from the initially generated synaptic matrix *S*, and became weaker for the trains obtained from the scaled matrices *S*_*α*_ = *α* · *S*. Specifically, the progressive dampening of the synaptic weights, achieved decreasing the values of *α* from 1 to 0 in steps of 0.2, resulted in values of CFI_MI_ gradually decreasing from about 0.8 to 0. While these trends were detected regardless of the parameter *b* for values of the synaptic strength ranging from high to intermediate (*α* = 1, 0.8, 0.6, 0.4), some inconsistencies were observed for *b* = 4 and *b* = 5 in the case of very low coupling strength (*α* = 0.2) or complete lack of synapses (*α* = 0). In these cases CFI_MI_ cannot detect absence of correlation; moreover in the case of *b* = 5, CFI_MI_ = 1. This behaviour is related to the fact that for low values of *α* both excitatory and inhibitory cells exhibit almost constant FR. In such cases, imposing a large idle threshold (large *b*) diminish the alternations between the two binary states (working/non-working); for *b* = 5 each neural unit remains in the working mode throughout the whole simulation time and the unitary profiles for both units prevent reliable MI analysis (CFI_MI_ = 1). Therefore, lower values of the input parameter for CFI_MI_ analysis are recommended: the choice of *b* = 2 or *b* = 3 allows for an adequate idle threshold estimation, such that the index exhibits limited variability and is able to follow the decrease in coupling strength as well as its absence in the cases of negligible or null synaptic weight (*α* = 0.2, *α* = 0). As thoroughly examined in (Mijatović et al., 2018), the value of idle threshold relates to the neuron’s ISI distribution to reflect a meaningful periods of neuron’s quiescence.

**Fig. 6.**
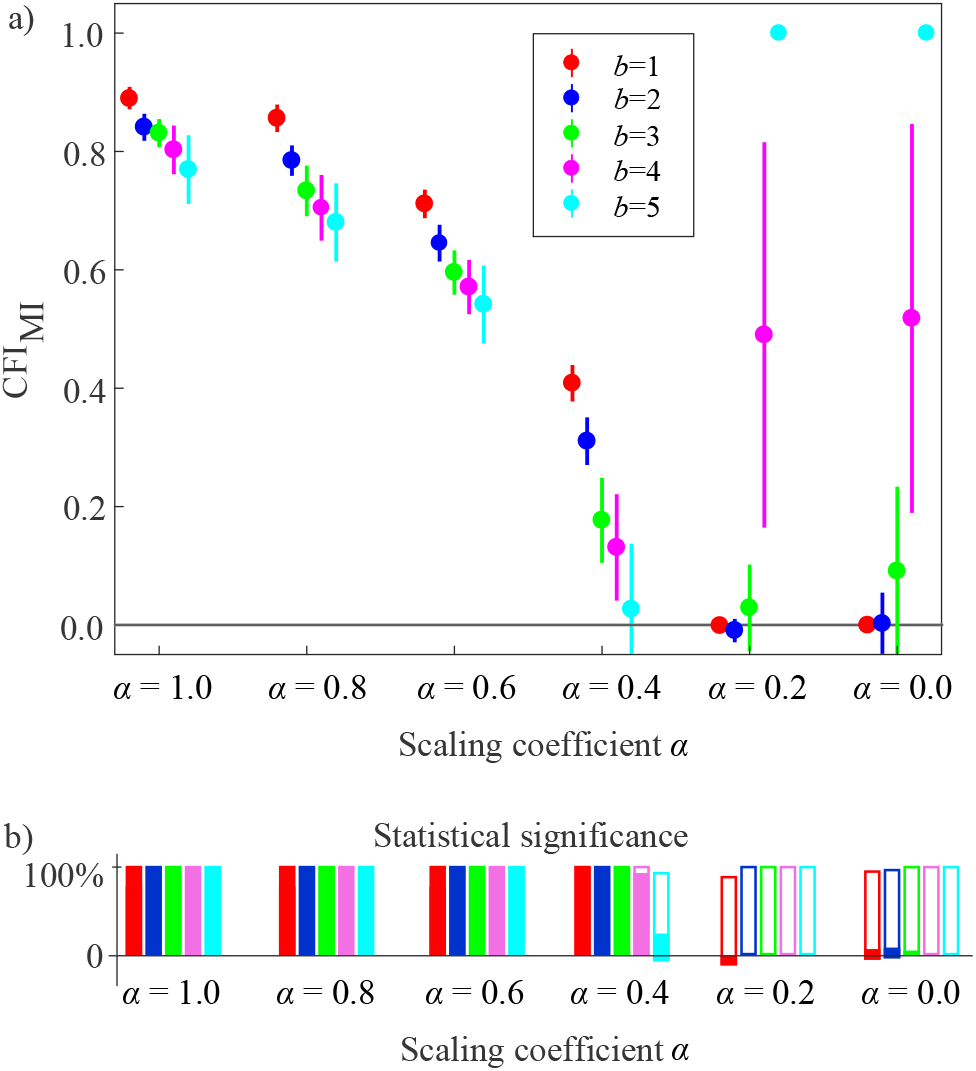
Simulation of RS-LTS neural networks. a) Distribution of the CFI_MI_ estimates (mean ± standard deviation over 100 pairs of neural responses) computed as a function of the scaling coefficient *α* applied to the initially generated synaptic matrix *S* for different values of the input parameter *b*; percentage of neural response pairs exhibiting significant positive correlation (filled bars above zero), absence of correlation (unfilled bars) or significant anti-correlation (filled bars below zero).

Fig. 7 illustrates with more detail the analysis conducted on the realistic network model. The pseudocolor representation of the full synaptic matrices (1000×1000 cells) of the simulated RS-LTS network is reported in Fig. 7a), showing how such matrices are modulated by the scaling coefficient *α*. It is worth noting that the strength of synapses conducting the outputs from excitatory cells is positive, while that corresponding to inhibitory cells is negative. The sub-matrices (10×10) in Fig. 7b) map the values of the CFI_MI_ index computed with input parameter *b* = 2 on the trains from the randomly selected RS and LTS units. Representative time windows of the activity of the selected cells are reported in Fig. 7c) to aid interpretation. Results confirm that the gradual weakening of the synaptic strength leads to a progressive decrease of the CFI_MI_ values. The raster plots confirm the presence of a fully correlated activity involving all cells for *α* = 1, which becomes progressively less synchronized in time as *α* decreases; when *α* = 0.2, the activity is sparse and evidently uncoupled due to the prevalence of the noisy thalamic inputs over the weak synaptic strengths. As a consequence, the CFI_MI_ index takes values around 0.82, 0.8, 0.63, 0.34 and 0 as seen in the color-coded distributions of Fig. 7b) and also in Fig. 6a) for *b* = 2. The rather uniform values of CFI_MI_ inside each matrix of Fig. 7b) can be explained by the similar spike timing behavior observed for any pair of cells in Fig. 7c).

**Fig. 7.**
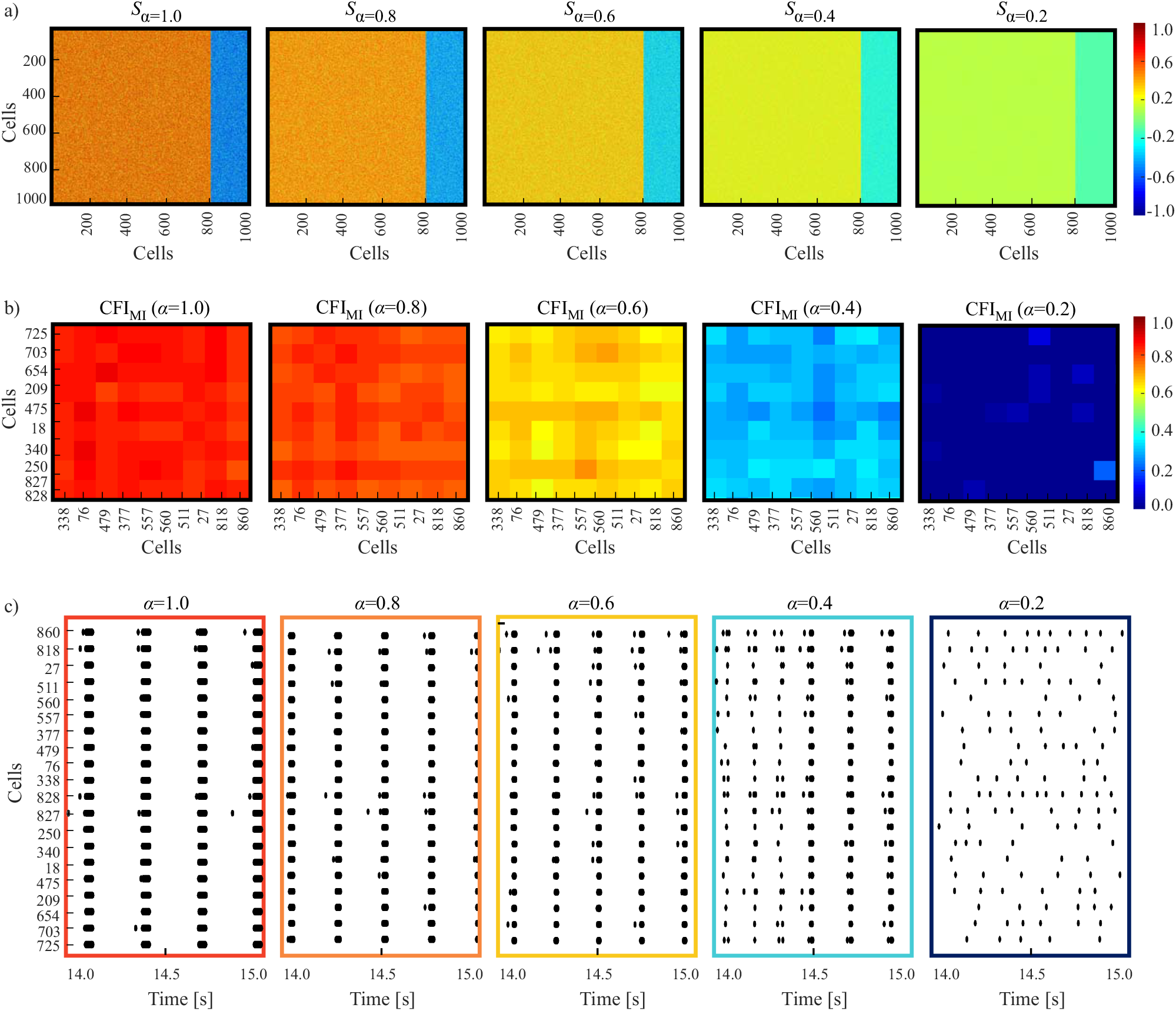
CFI_MI_ analysis of simulated RS-LTS neural networks. a) Synaptic matrices *S*_*α*_ locating the links between 1000 RS-LTS randomly connected cells, obtained scaling the initially generated synaptic matrix *S* by the coefficient *α* ∈ {1.0, 0.8, 0.6, 0.4, 0.2}; b) CFI_MI_ estimated for a randomly selected 10×10 RS-LTS sub-network (including both excitatory and inhibitory cells); c) one-second time-frame of spiking activity of the twenty randomly selected cells, which are fed with different synaptic inputs in terms of their strengths as shown in a). The parameter *b* is set to 2. The case *α* = 0 is not shown as it produced the same results as the case *α* = 0.2.

### 3.2 Application to experimental data

The ability of the proposed CFI_MI_ index to identify correlated firing activity in experimental data was evaluated using extracellular recordings by MEA across the population of RGCs, as described in Sect. 2.4. In particular, we have examined the extent of concurrency in firing activity for pairs of RGCs from the same retina exposed to the gradual increase in ambient light with superimposed white noise. First, the binary profiles representing the recorded firing activity of each cell were generated for the whole recording (with parameter *b* = 3). Then, after dividing the total recording time into 20 non-overlapping segments of approximately 45 s each, the CFI_MI_ index was estimated between spike train pairs over each temporal segment. This segmentation was motivated by the interest in studying the evolution of neural synchrony over time, producing time-resolved maps of the concurrent firing activity during periods of gradually increasing illumination of the photosensitive RGCs. The resulting matrices of pairwise CFI_MI_ of all RGCs in each retina are not illustrated as their size surpasses the discernible level of detail. To exemplify the informative value of CFI_MI_ we present the CFI_MI_ sub-matrix for the subset of 11 RGCs from one of the retinal explants, characterized by the same type of response (OFF) and significant synchronized activity. This is one of the subsets uncovered by distinctive CFI_MI_ patterns which indicated their tendency to fire nearly simultaneously more often than expected by chance. The synchronized activity is depicted in Fig. 8. Each row of the CFI_MI_ matrix in Fig. 8a) depicts the values of the index computed over the 20 temporal segments for one pair of cells; there are 55 rows in this matrix as 11 cells result in 55 pairs when symmetry is taken into account and pairs comprising same cells are excluded. Fig. 8b) reports the corresponding matrix of statistical significance for the CFI_MI_ index, coded with three colors (green: non-significant; brown: significant positive correlation; blue: significant anti-correlation). The figure shows an alternation between periods of uncoupled activity and periods with clear prevalence of synchronized firing patterns (e.g., time segments 7-8, 13, 20, see also raster plots in Fig. 8c).

**Fig. 8.**
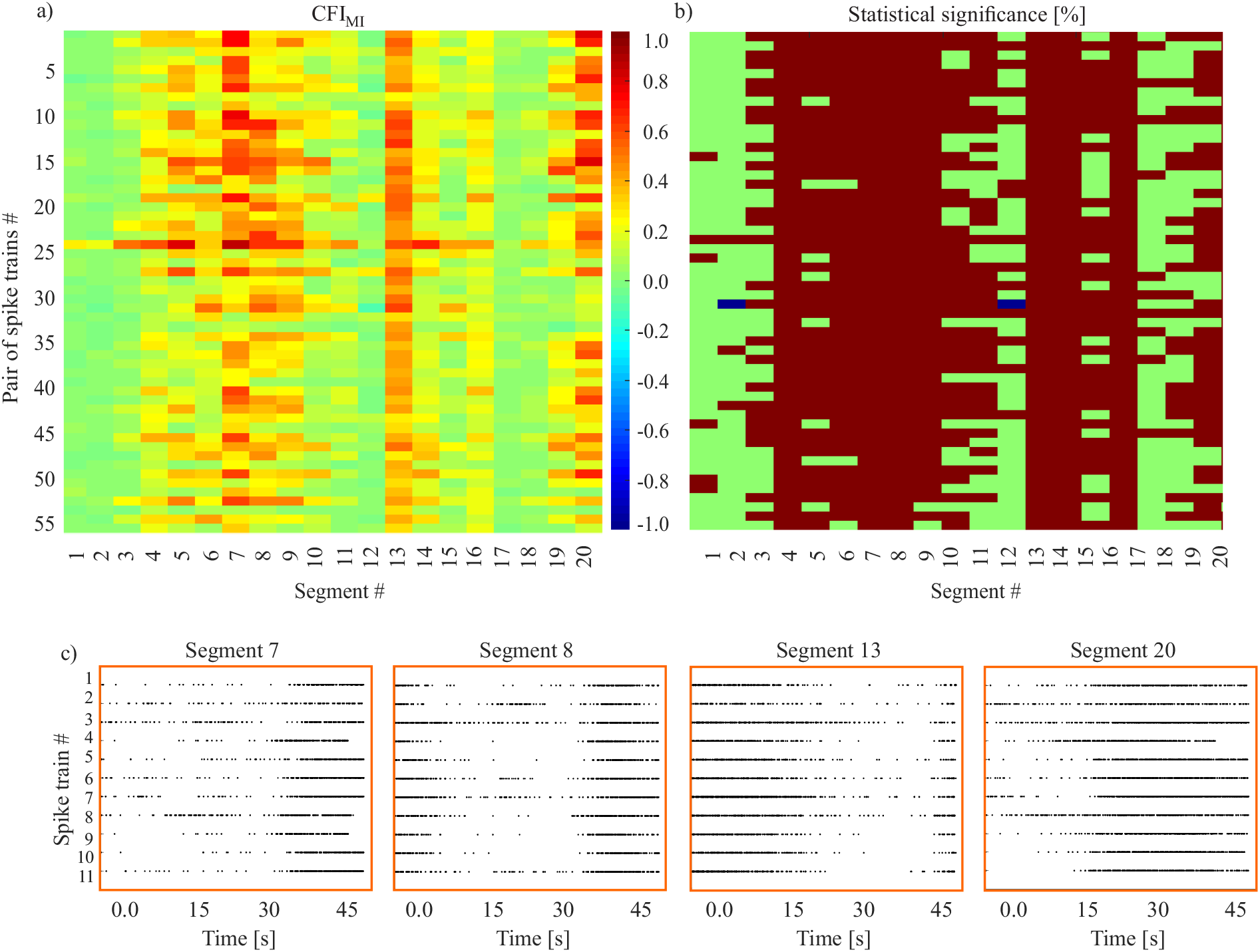
Computation of CFI_MI_ on an exemplary subset of the experimental data set. a) Matrix of CFI_MI_ estimated in a subpopulation of mostly adjacent 11 OFF cells (resulting in different 55 pairs of coupling values - matrix rows) for 20 temporally aligned segments (columns); b) corresponding matrix of statistical significance (blue: significant anti-correlation; green: non-significant correlation; brown: significant positive correlation); c) raster plots of four selected segments of the 11 ipRGc spike trains associated with high and statistically significant coupling. The CFI_MI_ input parameter *b* is set to 3; similarly results can be achieved for *b* = 2.

## 4 Discussion

We have described and evaluated a new approach to characterize the tendency of two neurons to exhibit temporally coordinated firing activity. The approach quantifies in a pairwise fashion the extent of concurrent firing activity between the neural responses, measured as the synchronization of the state transitions occurring in two simultaneously recorded neural spike trains. To do this, we apply information-theoretic metrics to the binary profiles of the two trains unfolded in time, obtained encoding the ISI intervals of each train into working and non-working states. Such coarse description is typical of symbolic computation, a popular approach whereby experimental signals are transformed into sequences of discretized symbols which retain essential information about the generating process (Porta et al., 2015; Daw et al., 2003). Symbolization is ubiquitously used in many fields of science ranging from geophysics and astrophysics to chemistry, fluidics, artifical intelligence and data mining (Daw et al., 2003). Applications in physiology and neuroscience are also popular (Porta et al., 2015), including those aiming specifically to describe event time series (e.g., cardiac heartbeat (Guzzetti et al., 2005) or neural inter-spike series (Steuer et al., 2001). In this work, we have exploited the advantages of the symbolic representation (e.g., robustness to noise, simplified representation of system’ states, reduction of redundancy, smoothing of non-stationarity, computational efficiency) in a bivariate fashion, proposing a MI measure able to quantify concurrency in the alternation between firing and quiescence periods of the spiking activity of two neurons. Additionally, the introduced modification to MI improves the definition of our CFI_MI_ index to ensure its clear interpretation as a standard correlation measure (range [−1,1]), where the sign indicates the type of correlation.

The proposed measure is assumption-free with respect to the ISI distribution of a spike train, and requires the setting of a single parameter to achieve a binary representation of the firing activity. It is worth noting that classification of ISIs into binary states is not guided by a threshold common to both trains, neither follows a one-fits-all principle. The threshold for detection of periods of attenuated activity (here named “non-working”) relates to the firing properties of each neuron and to the corresponding ISI distribution: *thr*_**I**_ = *b* · *mean*(ISI stream). Selection of the parameter *b* was examined in a previous study where the value *b* = 3 was adopted and further used to obtain the unique probabilistic characterization of the firing activity for each neuron (Mijatović et al., 2018). Here, we addressed the dependence of CFI_MI_ on the selection of this parameter in several simulation scenarios, exploring the alternative values *b* ∈ {1, 2, 3, 4, 5}. The values 2 or 3 resulted in a stable CFI_MI_ estimates, reflecting appropriately the ground truth set for the correlations between spike train pairs. Interestingly, selection of the value *b* = 3 is in line with the research on the cluster isolation criterion (Fred and Leitão, 2003), where it was shown that increments of a dissimilarity measure between neighboring patterns have exponential distribution regardless of the initial data generation model. The theoretical analysis of exponential distributions provided in (Fred and Leitão, 2003) has indicated that selecting the threshold for cluster isolation as a multiple of the mean value of an exponential distribution covers most of the distributions when a multiple takes values from the range [3, 5]. As the ISI distribution exhibits an exponential decay for long ISIs regardless of the strength of the fluctuating input (Ostojic, 2011), the suggested idle threshold as *b* · *mean*(ISI stream) can efficiently separate ISIs of short or moderate duration from longer ISIs.

Compared with existing pairwise measures of neural synchrony (Cutts and Eglen, 2014), the proposed CFI_MI_ index is different in the fact that, focusing on the co-occurrence of working and non-working states in two spike trains, it disregards the fine time-resolution properties of neural firing. In this way, at the price of losing information, our measure is robust against noise and free from the setting of resolution parameters. Nevertheless, as it is ultimately designed to capture correlations in neural spike trains, a comparison with existing measures is envisaged. In a preliminary investigation (Mijatovic et al., 2020, accepted paper), we used synthetic Poisson trains to compare our index with three well-established correlation measures: the Spike Count Correlation Coefficient (SCCC) (Eggermont, 2010), Kerschensteiner and Wong correlation (KWC) (Kerschensteiner and Wong, 2008), and the Spike Time Tiling Coefficient (STTC) (Cutts and Eglen, 2014) with respect to correlation range and robustness to the firing rate and duration of recording. Besides to higher robustness to variations in the firing rate and duration of recording, CFI_MI_ exhibits smaller variance in the detection of uncoupled firing activities and higher sensitivity to variations in coupling (Mijatovic et al., 2020, accepted paper). However, the interpretation of the CFI_MI_ values and relation to the other established synchrony (correlation) measures have to be elaborated in terms of both underlying methodology and temporal scale.

The proposed measure satisfies all the necessary properties formalized in (Cutts and Eglen, 2014) for a measure of neural synchrony: symmetry, boundedness, robustness to both firing rate and recording duration, and stability with respect to small variations in the input parameters. These properties were tested both theoretically and in controlled simulation scenarios involving coupled and uncoupled Poisson spike trains. In a theoretical example, we showed that CFI_MI_ is strictly bounded and can reflect either positive correlation, absence of correlation or anti-correlation, taking values equal to 1, 0 and −1, respectively when the spike trains produce identical, independent and opposite binary profiles.

In simulations of uncoupled Poisson trains we showed the stability of the measure across different firing rate ratios and duration of the two trains. In a customized simulation of coupled Poisson spike trains enabling fine tuning of correlation and anti-correlation levels between spike trains, we investigated the various levels of sensitivity to positive and negative correlations; exploring different values of the modulation parameter *γ* covering the whole correlation range, and different values of input parameter *b* in [1, 5], we showed that the sign of CFI_MI_ unequivocally corresponds to the sign of correlation between the firing patterns.

The simulation of a fully connected neural network of RL-LTS units provided a more complex but realistic scenario, inspired by the mammalian cortex. In this scenario, we assessed the capability of the CFI_MI_ index to reflect the changes in a coupling strength between neural units as defined by the synaptic matrix *S* (Sect. 3.1). The linear scaling of the coefficient in *S* led to a progressive decrease of the synaptic strength and weakening of the neural inputs, and consequently to reduced and decoupled overall firing activity. The CFI_MI_ index scaled with the coupling strength, ranging from highly correlated to non significant values. Its sign did not reflect the type of synapse (excitatory and inhibitory, i.e. positive and negative); this is however expected because, as long as the connected neural units exhibit synchronous concomitant firing as a result of the synaptic connections, the index should reflect the strictly positive correlations.

We also provide evidence for the applicability of the proposed approach to real spike train data, showing how it can uncover concomitant firing patterns in experimental recordings of retinal ganglion cells of mouse retinal explants. The group of 11 RGCs from the same retina, exemplified in Sect. 3.2, was identified by the specific pattern of the pairwise CFI_MI_ index values that revealed highly synchronized segments of their activation. This result is consistent with the previous research, extensively documented in many species, showing that RGCs frequently exhibit synchronized firing reflecting functional connectivity (Schnitzer and Meister, 2003; Ishikane et al., 2005; Shlens et al., 2008). Synchronized firing patterns can include at least 10 RGCs (Schneidman et al., 2006; Shlens et al., 2006), although their full extent is unknown (Shlens et al., 2008). Previous characterization of the patterns of electrical activity in primate RGCs reports that the structure of synchronized patterns is determined by the cell type, and may be understood almost entirely based on pairwise interactions mostly limited to adjacent cells in the neural mosaic (Shlens et al., 2006). In this context, we identified a small group of synchronized cells by means of their CFI_MI_ values, which resulted as statistically significant according to the hypothesis test employed. The identified synchronized cells exhibit temporal patterns of concurrent firing that alternate intermittently during the exposure to gradually increasing illumination. The absence of a clear dependence of the CFI_MI_ value on the gradual change in illumination might suggest that synchronized activity is not necessarily related only to the absolute value of stimulus intensity, just as their firing rate is, as shown in (Milosavljevic et al., 2018). However, the pattern of co-occurrence of the firing activity in the explored sub-population (Fig. 8a) reveals more coherent activation as the illumination level increases, further confirmed by the raster plots (Fig. 8c). This implies that not only the values of CFI_MI_, but also its rate and the pattern of its temporal changes, should be used to explore the response of synchronized firing activation to a stimulus presentation, or to any other experimental maneuver.

In conclusion, the proposed information-theoretic approach provides a simple and efficient way for the detection of synchronously firing units, yielding an index whose value and sign indicate the preference of two neurons to fire simultaneously or in alternation. Future work is envisaged to extend the sensitivity of the proposed measure to more than two functional states, e.g. differentiating bursting from moderate firing in the detection of coupled activity; further research will also point to the description of multivariate and dynamic interactions for allowing the causal analysis of networked neural interactions. These extensions will facilitate the characterization of multivariate correlations in a large variety of experimental settings and neurophysiological states.

## Acknowledgments

This research has been supported by the Ministry of Education, Science and Technological Development through the project no. 451-03-68/2020-14/200156: “Innovative scientific and artistic research from the FTS (activity) domain” and from the European Union’s Horizon 2020 research and innovation programme under Grant Agreement number 856967. Luca Faes acknowledges funding from Ministero dell’Istruzione, dell’Universitàe della Ricerca—PRIN 2017 (PRJ-0167), “Stochastic forecasting in complex systems”. The authors express gratitude to Leonardo Ricci and Alessio Perinelli for sharing Matlab code for JODI method, to Nina Milosavljevic for proving experimental data set, and to Danica Despotovic for very useful discussions.

## Information Sharing Statement

The software implementation of the proposed approach, including the algorithms for binary state representation of the spiking activity (Mijatović et al., 2018), computation of CFI_MI_ index, and assessment of its statistical significance based on surrogate data analysis (Ricci et al., 2019), are collected in the CFI MI MATLAB toolbox, which can be freely downloaded from https://github.com/mijatovicg/CFI-MI.

